# Toward pharmacologic therapy for glioblastoma: Identifying inhibitors of very long-chain acyl-CoA synthetase 3 (ACSVL3)

**DOI:** 10.1101/2025.07.02.662811

**Authors:** Emily M. Clay, Xiaohai Shi, Elizabeth A. Kolar, Mayur Mody, John E. Locke, Paul A. Watkins

## Abstract

Brain tumors, in particular glioblastoma multiforme (GBM), are among the most aggressive and difficult to treat human neoplasms. Even with combined surgery, radiation and chemotherapy, the 5-year survival rate for GBM is only ∼7%. Thus, new treatment approaches are needed. We previously found that the fatty acid metabolism enzyme “very long-chain acyl-CoA synthetase 3” (ACSVL3) is overproduced in human glioma tissue and in glioblastoma cell lines such as U87MG cells. These cells exhibited malignant growth properties in culture and were tumorigenic in nude mice. When either knockdown or knockout strategies were used to deplete U87MG cells of ACSVL3, they adopted a more normal growth rate and produced significantly fewer, slower growing tumors in mice. An inhibitor of ACSVL3, if identified, could prove to be a valuable pharmacotherapeutic agent in GBM. Therefore, we sought to identify small molecule compounds that decrease or block the enzyme activity of ACSVL3, as measured by the formation of stearoyl-CoA from the 18-carbon saturated fatty acid stearic acid, a preferred substrate for ACSVL3. We approached this in two ways. First, we tested several compounds that were previously shown to inhibit the activity of a structurally and functionally related enzyme, ACSVL1. Several compounds tested showed inhibition of stearoyl-CoA formation in U87MG cells when added to an in vitro enzyme assay. These included drugs triflupromazine, phenazopyridine, chlorpromazine, emodin, and perphenazine which are approved for treating other conditions. Also inhibitory to stearoyl-CoA production were several compounds from a ChemBridge Corporation library designated CB2, CB5, CB6 and CB 16.2. One caveat regarding interpretation of these results is that in addition to ACSVL3, all cells including U87MG contain other acyl-CoA synthetases capable of using stearic acid as substrate. Therefore, we also measured stearoyl-CoA synthetase activity in ACSVL3-deficient U87MG cells (U87-KO). If a drug or compound is an ACSVL3 inhibitor, it should decrease total conversion of stearate to stearoyl-CoA more in U87MG than in U87-KO cells. By this criterion, most of the tested compounds showed some ACSVL3-specific inhibition. At the screening concentration of 80μM drug, CB5 and CB16.2 showed the greatest potency to inhibit ACSVL3 enzyme activity; at 10 μM, CB5 still showed significant inhibition but CB16.2 did not. We conclude that these compounds are worthy of further investigation as potential therapeutic agents in GBM, but additional drugs that have greater specificity and are effective at significantly lower concentrations must also be identified. Therefore, our second strategy was to develop a high-throughput library screening assay. For this, we took advantage of the fatty acid transport capability of some ACSVL family members. ACSVL1, when heterologously expressed in COS-1 cells, promotes cellular uptake of the fluorescent fatty acid analog C_1_-BODIPY-C_12_; in contrast, overexpressed ACSVL3 does not. We used a domain-swapping strategy to replace the N-terminal 210 amino acids of ACSVL3 with the N-terminal 100 amino acids of ACSVL1, producing ACSVL1/3. Unlike ACSVL3, ACSVL1/3 robustly promoted C_1_-BODIPY-C_12_ uptake while retaining the catalytically active C-terminus of ACSVL3. Most of the drugs and compounds that decreased stearoyl-CoA synthetase inhibition also inhibited C_1_-BODIPY-C_12_ uptake in a concentration-dependent manner. Catalytically defective ACSVL1/3 mutants lost their ability to promote C_1_-BODIPY-C_12_ uptake. Thus, we conclude that chimeric ACSVL1/3 gained the fatty acid transport function of ACSVL1 while retaining the catalytic properties of ACSVL3. A pilot screening study of >1280 drugs from an approved drug library and >880 compounds from a library of drugs predicted to cross the blood-brain barrier detected more than 50 molecules that lowered C_1_-BODIPY-C_12_ by more than 3 standard deviations. Although secondary screening will likely exclude many or all of these, our findings support the notion that we have developed a viable method for detecting potential ACSVL3 inhibitors. Further characterization may reveal candidate pharmacologic agents for treatment of GBM and other cancers.

## INTRODUCTION

Glioblastoma multiforme (GBM) and other brain tumors are notoriously aggressive neoplasms, causing significant mortality and morbidity [Louis *et al*., 2016; Ostrom *et al*., 2016]. GBM is particularly resistant to treatment; despite current standard-of-care clinical management with surgery, radiation therapy and chemotherapy, the 5-year survival rate for GBM is only ∼7%. New approaches to treatment are thus sorely needed. Altered lipid metabolism has been frequently observed in many cancers [Bian et al., 2021; Broadfield et al., 2021], including changes in either expression or activity of several fatty acyl-CoA synthetases (ACS) [Cui et al., 2014; Hu et al., 2008; Wu et al., 2013; Gassler et al., 2013; Yamashita et al., 2000; Yoshii et al., 2009, Schug et al., 2015].

The ACSs are a group of enzymes that catalyze the conversion of fatty acids (FA) to FA-CoA. This activation reaction is required for FA to participate in nearly all downstream metabolic processes, including energy production via β-oxidation, remodeling of the FA via chain elongation or insertion/removal of double bonds, incorporation into complex lipids for energy storage or membrane synthesis, and post-translational covalent modification of proteins [Watkins, 1997]. The ACSs have been grouped into structurally-related families that roughly correspond to their preferred FA substrate chain length [Watkins et al. 2007]. Thus, the six enzymes of the “very long-chain”, or ACSVL, family are generally capable of activating FA whose carbon chains contain 20 or more carbons.

We previously reported that although ACSVL3 protein (gene name, SLC27A3) was not detected in adult mouse brain glial cells [Pei et al., 2004], it was highly overproduced in all human gliomas examined [Pei et al., 2009]. In addition to tumors, ACSVL3 was also found at high levels in glioblastoma cell lines such as U87MG and Mayo22 [Pei et al., 2009]. In culture, these cells exhibited malignant growth properties (lack of contact inhibition; adherence-independent growth) and produced rapidly growing subcutaneous and intracranial xenografts when injected into nude mice. When either knockdown [Pei et al., 2009] or knockout [Kolar et al., 2021] strategies were used to deplete U87MG cells of ACSVL3, they adopted a more normal growth rate in vitro and produced significantly fewer and slower growing tumors in mice. Thus, an inhibitor of ACSVL3, if identified, could prove to be a valuable pharmacotherapeutic agent in GBM.

To identify potential ACSLV3 inhibitors, we used a two-pronged approach. First, we asked whether inhibitors of homologous enzymes might also inhibit ACSVL3. The six members of the ACSVL family were independently identified and studied as “FA transport proteins” (FATP) [Black et al., 2009; Stahl 2004]. Schaffer and Lodish [1994] originally identified a protein that, when overexpressed in COS-1 cells, promoted uptake of the fluorescent FA analog C_1_-BODIPY-C_12_ and was named FATP (now FATP1 or SLC27A1). Subsequent investigation identified five additional homologs of FATP1 (FATP2-6) [Stahl, 2004]. Soon thereafter, sequence analysis revealed that ACSVL and FATP family members were identical proteins. However, although all six family members have demonstrable ACS activity, only FATP1, FATP2, and FATP4 showed significant transport capability when C_1_-BODIPY-C_12_ uptake was measured [Black et al., 2009; Sandoval et al., 2008]. Black, DiRusso and colleagues exploited the FA transport functionality to identify inhibitors of FATP2 (ACSVL1; SLC27A2) [Li et al., 2008; Sandoval et al., 2010]. In this study, we show that several inhibitors of ACSVL1 also inhibit ACSVL3.

Our second approach was to develop a similar fluorescence-based assay to screen chemical libraries for inhibitors of ACSVL3. Because ACSVL3 (FATP3) does not promote C_1_-BODIPY-C_12_ uptake, we employed a domain-swapping strategy to replace the N-terminal sequence of ACSVL3, which does not support FA transport, with the homologous domain from ACSVL1. Hybrid ACSVL1/3 constructs were found to facilitate uptake of fluorescent FA while retaining the catalytic domains of ACSVL3, and we used this as a high throughput library screening tool. We report here the identification of several molecules that inhibit ACSVL3 enzyme activity. We conclude that investigation of these drugs for their potential use as pharmacotherapeutic agents in glioma is warranted.

## MATERIALS AND METHODS

### Materials and general methods

Cell culture reagents were from CellGro Technologies except for fetal bovine serum (FBS) which was from Biosource International. [1-^14^C]Stearic acid was from Moravek, Inc. DMSO and C_1_-BODIPY-C_12_ were from Thermo Fisher Scientific. The Johns Hopkins Clinical Compound Library 20 was the kind gift of Dr. Jun O. Liu (Department of Pharmacology and Experimental Therapeutics, Johns Hopkins University School of Medicine). The ChemBridge CNS-set was obtained from the ChemBridge Online Chemical Store (hit2lead.com). Nucleotide sequences of all constructs were verified by the Johns Hopkins University Biosynthesis and Sequencing Core Facility. Cell authentication for U87MG cells was done within a year of study by the Johns Hopkins Genetics Resources Core Facility by short tandem repeat analysis using the PowerPlex 16 HS kit (Promega). Protein was measured by the method of Lowry et al. [1951] unless otherwise indicated.

### Cell culture

U87MG and COS-1 cells were obtained from American Type Culture Collection. The ACSVL3-knockout (U87-KO) cell line was produced with the CompoZr Knockout Zinc Finger Nuclease kit (Millipore Sigma) and characterized as described previously [Kolar et al., 2021]. Both lines were cultured in Dulbecco’s Modified Eagle’s Medium (DMEM) with 10% FBS. All cell lines were maintained at 37°C and 5% CO2 atmosphere.

### Drugs and compounds

Table 1 lists the common and systematic names of the drugs and compounds used in this study. Triflupromazine, phenazopyridine, chlorpromazine, emodin, and perphenazine were from Millipore Sigma. CB2, CB5, CB6 and CB16.2 were provided by Drs. Paul Black and Concetta DiRusso (Department of Biochemistry, Univ. of Nebraska, Lincoln NE). All were prepared as 1000x stock solutions in DMSO and added to cultures and assays to the final concentration indicated in each experiment. The final DMSO concentration was 0.1%.

**Table 1.**
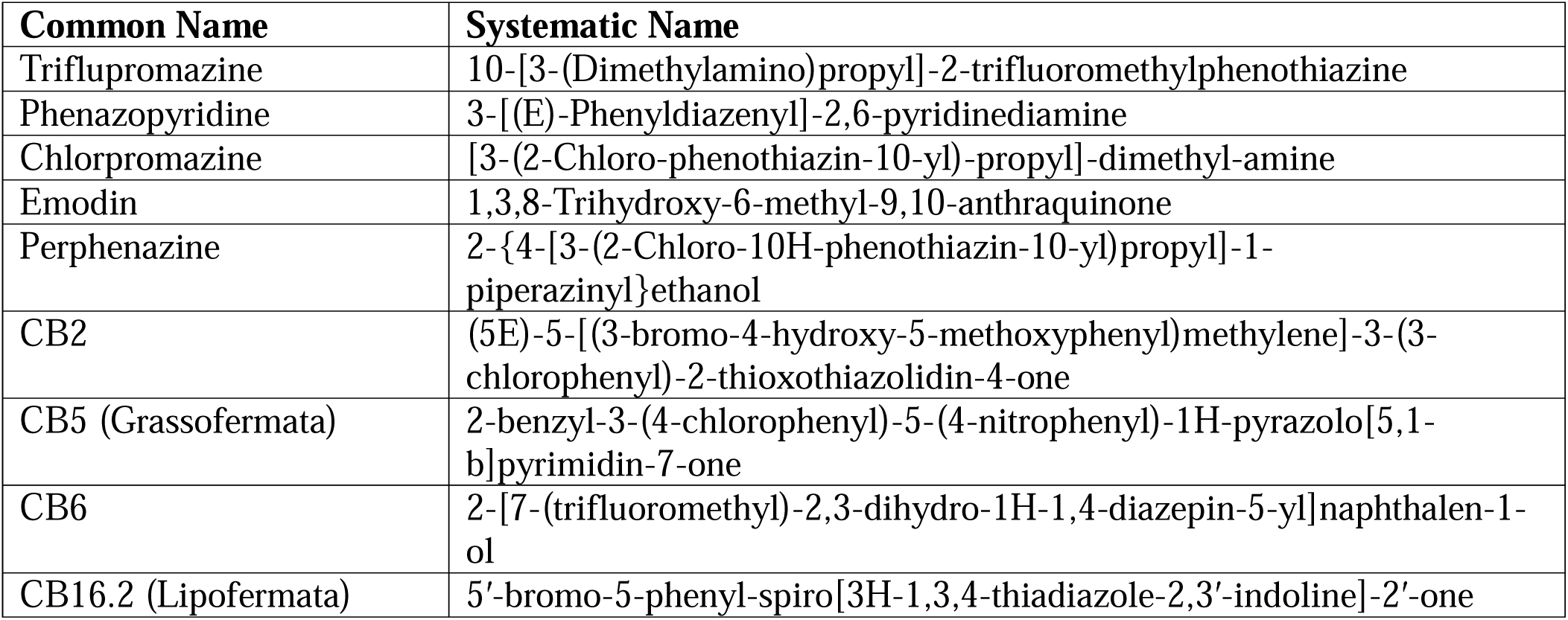
Drugs and compounds included in assays. The common name and systematic name of drugs and compounds used in this study are shown.

| Common Name | Systematic Name |
| --- | --- |
| Triflupromazine | 10-[3-(Dimethylamino)propyl]-2-trifluoromethylphenothiazine |
| Phenazopyridine | 3-[(E)-Phenyldiazenyl]-2,6-pyridinediamine |
| Chlorpromazine | [3-(2-Chloro-phenothiazin-10-yl)-propyl]-dimethyl-amine |
| Emodin | 1,3,8-Trihydroxy-6-methyl-9,10-anthraquinone |
| Perphenazine | 2-{4-[3-(2-Chloro-10H-phenothiazin-10-yl)propyl]-1-piperazinyl}ethanol |
| CB2 | (5E)-5-[(3-bromo-4-hydroxy-5-methoxyphenyl)methylene]-3-(3-chlorophenyl)-2-thioxothiazolidin-4-one |
| CB5 (Grassofermata) | 2-benzyl-3-(4-chlorophenyl)-5-(4-nitrophenyl)-1H-pyrazolo[5,1-b]pyrimidin-7-one |
| CB6 | 2-[7-(trifluoromethyl)-2,3-dihydro-1H-1,4-diazepin-5-yl]naphthalen-1-ol |
| CB16.2 (Lipofermata) | 5'-bromo-5-phenyl-spiro[3H-1,3,4-thiadiazole-2,3'-indoline]-2'-one |

### ACVL3 antibody

A monoclonal antibody to the C-terminus of human ACSVL3 was produced at the University of California Davis/NIH NeuroMab Facility (Project # FSAI183). A cDNA fragment encoding the C-terminal 175 amino acids of human ACSVL3 was cloned in frame into the EcoR1 and BamH1 sites of the pMalC2 expression vector. Transfection of this construct into E. coli strain BL21(DE3) and induction of protein synthesis using isopropyl β-D-1-thiogalactopyranoside (IPTG) resulted in production of a 62 kDa fusion protein containing Maltose Binding Protein (MBP) at the N-terminus and human ACSVL3 fragment at the C-terminus. The fusion protein was purified from lysed cells by affinity chromatography on Amylose resin (New England Biolabs) using the procedure of Reichelt et al. [2006] to remove endotoxins. After elution with 10 mM maltose, the fusion protein was cleaved using Factor Xa (Millipore Sigma). The liberated MBP portion (42.5 kDa) was removed by re-binding to the Amylose resin, leaving the ACSVL3 C-terminal fragment (19.6 kDa) in solution. This protein was concentrated using a 30,000 MWCO centrifugal concentrator (Amicon); dilution in maltose-free buffer and re-centrifugation was then used to remove maltose. After verifying its purity by gel electrophoresis, the ACSVL3 fragment was sent to NCI for antibody production. More than 100 hybridoma culture supernatants were subsequently received back from NCI and tested by western blot and immunofluorescence. Clones 1C1, 1E5, 2G2 and 3D11 proved to be excellent Mabs.

Immunofluorescence and western blotting were used to assess ACSVL3 expression in U87MG and U87-KO cells as previously described [Jia et al., 2007]. For both immunofluorescence and western blotting, the primary antibody was a 1:10 dilution of hybridoma clone 2G2 supernatant. The secondary antibody for immunofluorescence was goat-α-mouse Cy3 (Jackson ImmunoResearch; 1:150 dilution in PBS/0.01% BSA). For western blotting, the secondary antibody was goat α-mouse IgG-HRP conjugate (Santa Cruz Biotechnology; 1:8,000 dilution in 10% nonfat dried milk solution).

### Proteomic analysis of ACS expression

Proteomic analysis of U87MG cells and normal human astrocytes was performed as described previously [Kolar et al., 2021; Nirujogi et el., 2015; Kim et al., 2014]. Tryptic peptides were labeled with tandem mass tags, fractionated by reverse-phase liquid chromatography, and fractions analyzed using an Orbitrap Elite tandem mass spectrometer. The average of biological replicates was obtained and the relative expression of ACSs in astrocytes and U87MG cells was determined.

### Measurement of acyl-CoA synthetase activity

Cells grown in 10 cm culture dishes were treated with 1 mL of 0.25% trypsin to release cells from the plates. The cell suspension was pelleted by centrifugation and then washed twice with phosphate-buffered saline (PBS) containing a protease inhibitor cocktail (Roche). The pellets were resuspended in buffer containing 0.25 M sucrose, 1 mM Tris-HCl pH 7.5, and 1 mM EDTA (STE buffer). Acyl-CoA synthetase assays were essentially as described by Watkins et al. [1991]. Assays contained (final concentrations) 40 mM Tris-Cl (pH 8.0), 10 mM ATP, 1 mM MgCl_2_, 0.2 mM CoA, 0.2 mM dithiothreitol (DTT), 20 μM [1-^14^C]stearic acid (20,000 dpm/nmol; solubilized in 10 mg/ml α-cyclodextrin (Millipore Sigma) in 10 mM Tris pH 8.0), 30 μg cell protein, and drug or vehicle. Assay volume was 0.25 ml and the final volume of DMSO was 2 μl. After an incubation of 20 min at 37°C, ^14^C-labeled acyl-CoA products were separated from the [1-^14^C]stearic acid substrate by the method of Dole [1956] and quantitated using liquid scintillation counting. Results are presented as nmol stearic acid activated per 20 minutes per mg protein.

### Generation of domain-swapped ACSVL1/3

Sequence and ligation independent cloning (SLIC) as described by Li and Elledge [2012] was used to produce chimeric cDNA containing the N-terminal regulatory domain of ACSVL1 and the enzymatic C-terminal domain of ACSVL3. To accomplish this, nucleotides coding for amino acids 1-210 of ACSVL3 were replaced with those coding for amino acids 1-100 of ACSVL1. ACSVL1 and ACSVL3 were individually cloned into the mammalian expression vector pcDNA3 (Invitrogen) as previously described [Steinberg et al., 1999; Pei et al., 2004]. The N-terminal 300 bp fragment of ACSVL1 gene was PCR-amplified by Phusion Polymerase with primers 5’-cttggtaccgagctcGGATCCgccaccATGCTTTCCGCCATCTACAC-3’ (For-1, which contains a BamH1 restriction site) and 5’-acagttgctccaggtgacaggccgaggtggtcgtgcag-3’. The C-terminal 1,563 bp fragment of ACSVL3 gene was similarly PCR-amplified using primers 5’-CTGTCACCTGGAGCAACT-3’ and 5’-tgatggatatctgcagaattcTCAGATTCGAAGGTTTCCTG-3’ (Rev-1, which incorporates an EcoR1 site). 1 µg each of the pcDNA3 vector linearized with BamHI and EcoRI and the two inserts were separately treated with T4 DNA Polymerase (New England Biolabs). The reactions were stopped with 1/10 volume of 10 mM dCTP. Next, the vector and two inserts were annealed at 1:1:1 ratio with RecA protein (Epicentre Biotechnologies). 2 µl of the annealing reaction were used to electroporate E. coli DH10B competent cells, followed by plating and culture. Colonies containing the chimeric sequence were sequenced and amplified. After transfection into COS-1 cells using FuGENE6 (Promega), colonies resistant to G418 were selected. The resulting hybrid construct is referred to as ACSVL1/3.

### Site directed mutagenesis

Two catalytically defective ACSVL1/3 chimera cell lines were created using site-directed mutagenesis as previously described [Pei et al., 2005]. Overlap extension PCR [Ho et al. 1989] was used to replace amino acids in highly conserved domains (Motif I and Motif II, shown in red typeface in Fig. 3), that were previously identified as being critical for catalysis [Black et al., 1997]. To produce ACSVL1/3Mut1, the first amino acid in Motif I (bold red typeface in Fig. 3) was changed from Phe to Ala. Two overlapping fragments were amplified by PCR using full-length ACSVL1/3 as template and primers incorporating the alanine substitution (shown in bold underline). The first reaction used forward primer For-1 and reverse primer 5’-cccgtggtgccagaggt**<u>tgc</u>**gatgtacaggcacgtgt-3’ (Rev 2) and amplified a 689 bp fragment encoding the first 223 amino acids of ACSVL1/3. The second PCR reaction used forward primer 5’-acacgtgcctgtacatc**<u>gca</u>**acctctggcaccacggg-3’ (For-2) and reverse primer Rev-1 that amplified a 1211 bp fragment encoding the C-terminal 402 amino acids. After gel purification, the two overlapping PCR products were used as template for another round of PCR using For-1 and Rev-1 as forward and reverse primers, respectively. After cloning into the pcDNA3 vector using BamH1 and EcoR1, inserts containing the correct sequence were transfected into COS-1 cells. Clones stably expressing ACSVL1/3Mut1 were selected using G418.

The same procedure was used to change the Arg located at the center of Motif II (bold red typeface in Fig. 3) to Ala to yield ACSVL1/3Mut2, with the following exceptions. The first PCR used forward primer For-1 and reverse primer 5’-ggtgtctccagt**<u>agc</u>**atcatggaagcggagaaaaccttg-3’ (Rev-3) that amplified a 1452 bp fragment encoding the first 483 amino acids of ACSVL1/3. The second PCR reaction used forward primer 5’-caaggttttctccgcttccatgat**<u>gct</u>**actggagacacc-3’ (For-3) and reverse primer Rev-1 that amplified a 450 bp fragment encoding the C-terminal 149 amino acids.

### C_1_-BODIPY-C_12_ uptake in COS-1 cells expressing ACSVL1/3

Cloning vector pcDNA3 containing the full-length sequences of human ACSVL3 and ACSVL1 as well as the empty vector were transfected into COS-1 cells. Clones stably expressing these constructs were selected by G418 resistance for comparison to and characterization of chimeric ACSVL1/3. COS-1 cells stably expressing each construct were plated onto glass coverslips and allowed to attach in normal media. The media was aspirated and replaced with serum-free media for 30 minutes, and then incubated with 2 μM 4,4-difluoro-5-methyl-4-bora-3a,4a-diaza-s-indacene-3-dodecanoic acid (C_1_-BODIPY-C_12_) for 20 minutes. Coverslips were washed with PBS and visualized via fluorescence microscopy to assess C_1_-BODIPY-C_12_ uptake.

To quantify C_1_-BODIPY-C_12_ uptake, COS-1 cells stably expressing ACSVL3, ACSVL1, ACSVL1/3, and the empty vector pcDNA3 were plated into opaque-walled 96-well plates and allowed to grow to confluence under normal growth conditions. The culture media was removed and replaced with serum-free DMEM for serum starvation. After 1 hour, the serum-free media was replaced with DMEM containing 5 μM C_1_-BODIPY-C_12_, 5 μM bovine serum albumin (BSA) and 1.97 mM trypan blue. Inhibitors were added from 1000x stock solutions in DMSO. The plates were incubated for 20 minutes at 37°C and 5% CO2. The trypan blue quenches fluorescence of extracellular C_1_-BODIPY-C_12_ that is not taken up by cells. Fluorescence of C_1_-BODIPY-C_12_ internalized by cells was quantitated using a SpectraMax M5 microplate reader.

### Screening of chemical libraries

The assay described above was used to screen portions of two drug/chemical libraries. The Johns Hopkins Clinical Compound Library contains 2560 compounds in 96-well format [Chong et al., 2006]. Most are approved drugs but also present were natural products and drugs in clinical trials. 16 plates were screened. The ChemBridge CNS-MPO library contains 50,000 compounds in 96-well format, and 11 plates were screened. Plate columns 1 and 12 contained cells expressing either ACSVL1/3 or vector that were treated with vehicle (DMSO) alone; these were used to determine the upper (ACSVL1/3) and lower (vector only) C_1_-BODIPY-C_12_ uptake limits. These values were used to calculate Z-factor (Z′), defined as: Z′ = 1 − (3 × SSD/R), where R = absolute mean difference and SSD = sum of the standard deviations (SDs) [Zhang et al., 1999]. A Z-factor of 0.5–1.0 is excellent, while 0–0.5 is good and less than 0 is unacceptable.

## RESULTS

### Rationale for testing inhibitors of ACSVL1 for their ability to inhibit ACSVL3

In human brain, endogenous ACSVL3 is only weakly expressed. In contrast, our laboratory demonstrated that ACSVL3 was overexpressed in all of 79 human gliomas examined, as well as in several glioma cell lines including U87MG and Mayo-22 [Pei et al., 2009]. These cell lines exhibit rapid growth on culture dishes, are not contact-inhibited, form colonies when grown in suspension culture, exhibit abnormal Akt signaling and form tumors when injected either subcutaneously or orthotopically into nude mice [Pei et al., 2009]. Depletion of ACSVL3 in these cell lines by either siRNA-mediated knockdown [Pei et al., 2009] or zinc finger nuclease-mediated knockout [Kolar et al., 2021] significantly reversed these malignant growth properties. Knockdown or knockout cells had a more normal growth rate on culture dishes, exhibited contact inhibition and had reduced ability to grow in suspension culture. When injected subcutaneously or orthotopically in nude mice, fewer xenografts developed and those that did had a significantly reduced growth rates. In toto, our findings strongly suggested that ACSVL3 is a promising therapeutic target in glioma. While results of ACSVL3 knockdown or knockout in mice are promising, a more realistic approach to treating human cancer patients would be to identify a small molecule inhibitor of ACSVL3, i.e. a drug, that would have antineoplastic properties. To date, no such drug is known. One approaches to identify candidate ACSVL3 inhibitors is to test drugs that inhibit structurally and functionally related enzymes, e.g. other ACSVL family members. Black, DiRusso and colleagues identified several compounds that inhibit ACSVL1 (FATP2) [Li et al., 2008; Sandoval et al., 2010], and we hypothesized that one or more of these may also inhibit ACSVL3.

We verified that U87MG cells are an appropriate source of ACSVL3 for this study by immunofluorescence and western blotting using a monoclonal antibody produced by the NIH NeuroMAbs program (see Methods). Robust ACSVL3 expression in U87MG cells was detected using either indirect immunofluorescence or western blotting (Figure 1). Similarly, both methods verified that U87-KO cells lack ACSVL3 expression (Figure 1). We previously determined that ACSVL3 activated long- to very long-chain FAs; the highest specific activity of >50 nmol of FA activated to FA-CoA/20 min/mg protein was measured when radiolabeled stearate (C18:0) was the substrate [Kolar et al., 2021]. Therefore, we chose activation of [1-^14^C]stearate in U87MG cells as the optimal assay for testing the effect of potential inhibitors on ACSVL3 activity.

**Figure 1.**
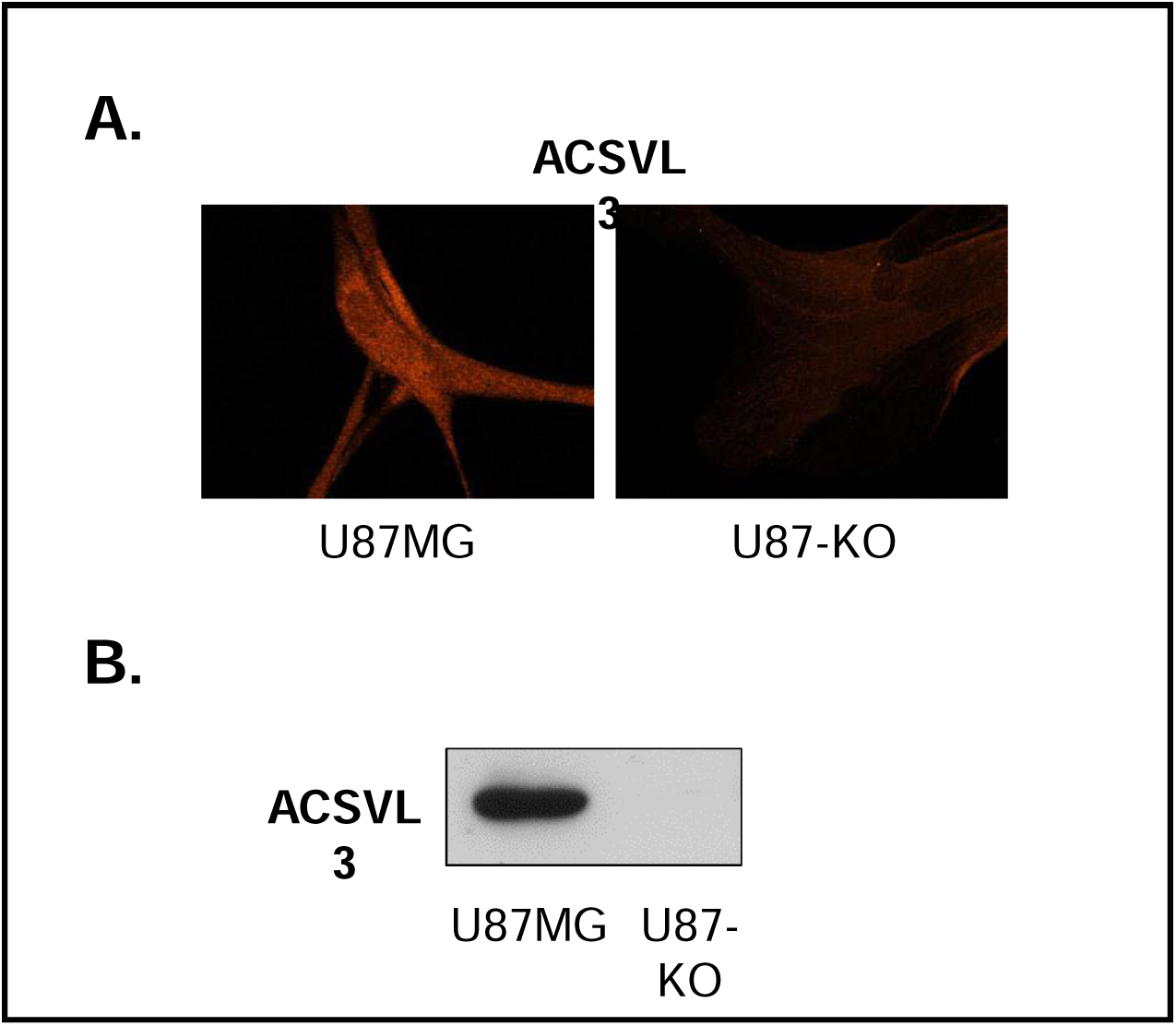
Expression of ACSVL3 in U87MG and U87-KO cells. A. Immnofluorescence. Cells were plated on glass coverslips and allowed to grow for 3 days. Cells were then fixed, permeabilized, incubated with human anti-ACSVL3 monoclonal antibody clone 2G2 and detected with Cy3-conjugated secondary antibody as described in Methods. B. Western blot. Cells were grown to near confluence and harvested by gentle trypsinization. 30 μg of protein from each cell type was subjected to SDS-PAGE and Western blotting as described in Methods. ACSVL3 was detected using monoclonal antibody clone 2G2, also as described in Methods. Both methods confirmed lack of ACSVL3 expression in U87-KO cells.

Fortuitously, U87MG cells proved to be a good cell line in which to test ACSVL1 inhibitors. Unbiased proteomic analysis revealed that while U87MG cells express ACSVL3, FATP1 and FATP4, expression of ACSVL1 (FATP2), FATP5 and FATP6 was below the level of detection by tandem mass spectrometry (Table 2). Thus, any inhibition of ACS activity observed in U87MG cells cannot be ascribed to effects on ACSVL1.

**Table 2.** Expression of ACS proteins capable of activating long- and very long-chain FA in normal human astrocytes and U87MG cells. Unbiased proteomic analysis of human astrocytes and U87MG cells was carried out as described in Methods. Protein expression in astrocytes is presented as intensity-based absolute quantification (iBAQ; column 1). The intensity of the highest expressed protein, ACSL3, was set to 100% and relative expression in astrocytes of the other ACSs calculated (column 2). Expression of ACSL3 in U87MG cells was then set to 100% and the relative expression of other ACSs determined (column 3). Both cell types express ACSs that are capable of activating long- to very long-chain FA and thus could theoretically use [1-^14^C]stearic acid as substrate. These include four ACSL family enzymes (ACSL1, 3, 4 and 5) and three ACSVL family enzymes (SLC274A1, 3 and 4). Other ACSs that can activate long- to very-long chain length FA, including ACSL6, SLC27A2, 5 and 6, and ACSBG1 and 2, were not detected (n.d.). Expression of ACSVL3 was about 10-fold higher in U87MG cells than in normal astrocytes, supporting our contention that ACSVL3 is overproduced in GBM cells and gliomas. ACSL5, which was not detected in astrocytes, was robustly expressed in U87MG cells. Notably, ACSVL1 was among the ACSs not detected in U87 cells, suggesting that this cell line is appropriate for testing whether inhibitors of ACSVL1 could inhibit ACSVL3.

| Gene | Other name(s) | Human astrocytes (iBAQ) | Human astrocytes (relative to ACSL3) | U87MG cells (relative to ACSL3) |
| --- | --- | --- | --- | --- |
| ACSL3 |  | 90.44 | (100) | (100) |
| SLC27A4 | ACSVL5, FATP4 | 55.62 | 61.5 | 94.3 |
| ACSL1 |  | 46.08 | 51.0 | 65.4 |
| SLC27A1 | ACSVL4, FATP1 | 36.32 | 40.2 | 32.4 |
| ACSL4 |  | 34.65 | 38.3 | 61.0 |
| SLC27A3 | ACSVL3, FATP3 | 3.3 | 3.6 | 38.4 |
| ACSL5 |  | (n.d.) | 0 | 50.5 |
| ACSL6 |  | (n.d.) | 0 | 0 |
| SLC27A2 | ACSVL1, FATP2 | (n.d.) | 0 | 0 |
| SLC27A5 | ACSVL6, FATP5 | (n.d.) | 0 | 0 |
| SLC27A6 | ACSVL2, FATP6 | (n.d.) | 0 | 0 |
| ACSBG1 | Bubblegum | (n.d.) | 0 | 0 |
| ACSBG2 | Bubblegum-related | (n.d.) | 0 | 0 |

### Several ACSVL1 inhibitors also inhibited ACSVL3

ACSVL1 inhibitors tested for their ability to inhibit ACSVL3 in U87MG cells included several approved drugs (Table 1, first five compounds). These were identified by Li et al. [2007] by high-throughput screening of the SpectrumPlus library (MicroSource) containing 2080 compounds. Also tested were four compounds identified by Sandoval et al. [2010] as inhibitors of ACSVL1 by screening a ChemBridge Corporation library containing 100,000 chemicals. When added to ACS assays at a concentration of 80 μM, all five drugs – triflupromazine, phenazopyridine, chlorpromazine, emodin, and perphenazine – inhibited total cellular ability to activate [1-^14^C]stearate (Figure 2A). The four ChemBridge ACSVL1 inhibitors – CB2, CB5, CB6 and CB16.2 – also inhibited activation of stearate to stearoyl-CoA by U87MG cells (Figure 2A).

**Figure 2:**
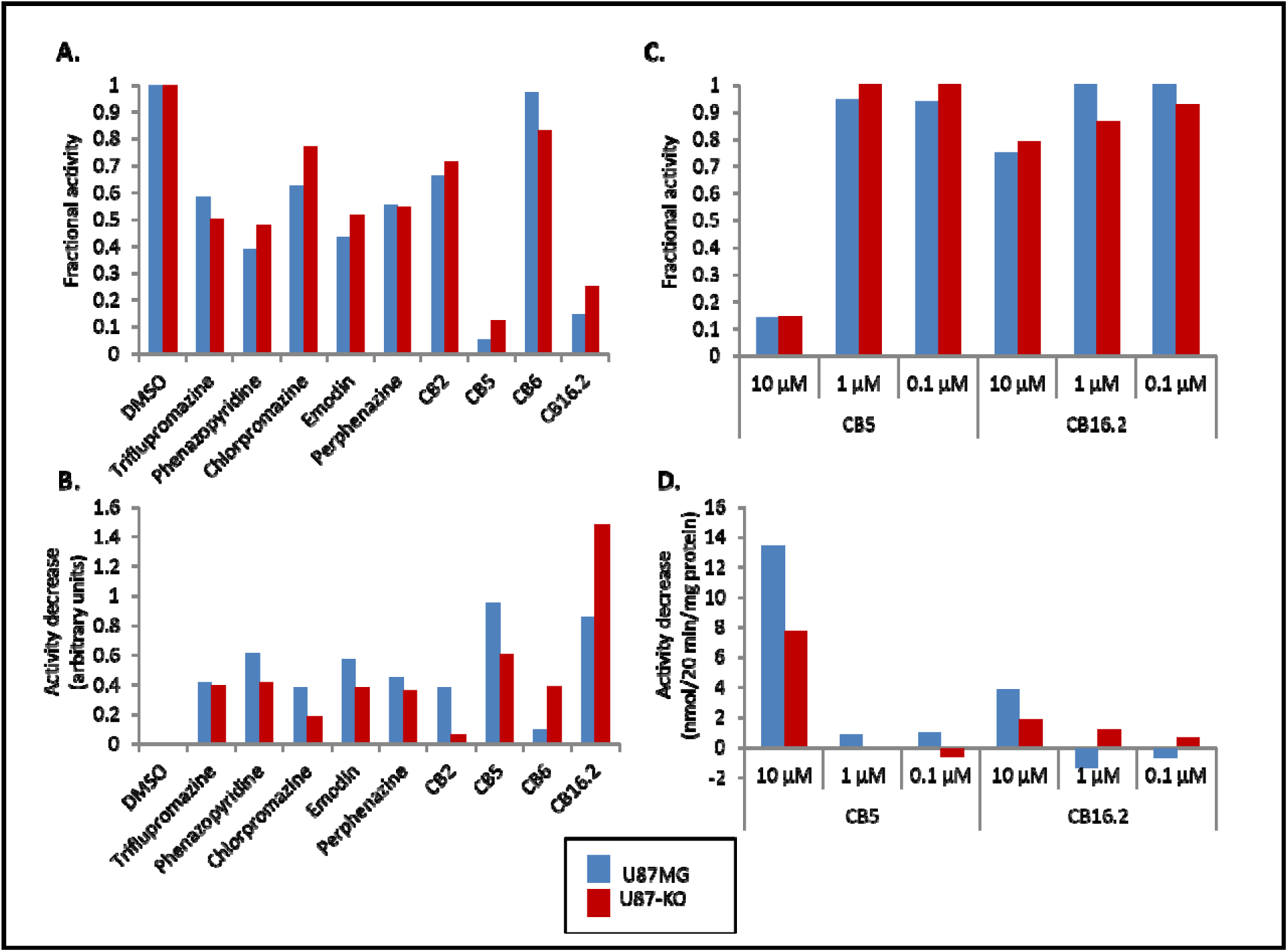
Inhibition of C18:0 activation by ACSVL1 inhibitors in U87MG and U87-KO cells. 80 µM of various ACSVL1 inhibitors was added to assays assessing the conversion of [1-^14^C]-labeled C18:0 to its CoA derivative in protein from both U87MG and U87-KO cells. For each drug or compound, the fraction of activity compared to assays containing DMSO (negative control) is shown (A). The total decrease in ACS activity, normalized to the DMSO control in each assay, is also shown (B). An inhibitor that is somewhat specific for ACSVL3 should exhibit either a higher fraction of activity in KO vs. WT, or a larger total difference in ACS activity in KO vs. WT. CB5 and CB16.2 were determined to be the most robust inhibitors and were added to ACS assays in smaller concentrations. Again, the fraction of activity (C) and the total decrease in ACS activity (D) are shown. 10 µM CB5 inhibited ACS activity robustly, 10 µM CB16.2 inhibited ACS activity partially (but less robustly than CB5), and lower concentrations of both drugs did not inhibit ACS activity.

An important caveat to the interpretation of these data is that cells, including U87MG, typically express several different ACS. As noted above (Table 2), six ACSs capable of activating long- and very long-chain FA are expressed in U87MG cells and the radioactive assay used measures the sum of activities of all ACSs present. For this reason, cellular capacity to activate [1-^14^C]stearate was also measured in U87-KO cells. Figure 2A shows the fraction of stearate activation in the presence of inhibitors compared to DMSO, while Figure 2B shows the difference in total conversion of stearate to stearoyl -CoA in the presence of the inhibitors. An inhibitor that lowers total conversion of stearate to stearoyl-CoA more in U87MG cells than in U87-KO cells would be considered to be somewhat specific for ACSVL3.

All of the ACSVL1 inhibitors except CB6 and CB16.2 lowered total conversion of stearate by a larger margin in U87MG cells compared to U87-KO cells. The following inhibitors showed a greater percent decrease in U87MG versus U87-KO cells: phenazopyridine, chlorpromazine, emodin, CB2, CB5, and CB16.2.

Since CB5 and CB16.2 strongly inhibited stearate activation at 80 μM, assays were repeated with lower doses of these two compounds. Again, the difference in inhibition in WT and ACSVL3-KO U87 cells was compared (Figure 2C and 2D). The inhibitory effect of CB16.2 was greatly reduced at these lower concentrations. CB5 was still significantly inhibitory at 10 μM, but not at 1 μM or 0.1 μM. Thus, CB5 shows the most promise as a potential ACSVL3 inhibitor. Further studies to determine whether CB5 abrogates the malignant behavior of U87MG cells are in progress.

### Development and characterization of a domain-swapped ACSVL1/3 hybrid enzyme

The identification by Black, DiRusso and colleagues of SpectrumPlus and ChemBridge library compounds as inhibitors of ACSVL1 exploited the dual functionality of certain ACSVL family members to facilitate the uptake of fluorescent or radiolabeled FAs as well as to catalyze the thioesterification of FAs to their CoA derivatives [Stahl 2004; Watkins 2008]. We wished to develop a similar assay to identify ACSVL3 inhibitors. However, unlike ACSVL1, ACSVL3 does not facilitate cellular uptake of FA as measured using the fluorescent FA C_1_-BODIPY-C_12_ [DiRusso et al. 2005]. Alignment of the amino acid sequences of ACSVL1 and ACSVL3 revealed that the C-terminal two-thirds of these proteins are more highly conserved than their N-terminal third (Figure 3). Since the C-termini contain several highly conserved motifs required for ACS enzyme activity [Watkins et al. 2007; Black et al. 1997], we hypothesized that perhaps the less well-conserved N-terminal domains have regulatory functions and/or contain domains necessary for FA transport. To address this possibility, we created a chimera that contains the N-terminus of ACSVL1 fused in-frame to the catalytic C-terminal domain of ACSVL3 (illustrated in Figure 4A). The hybrid construct was cloned into the mammalian expression vector pcDNA3 and transfected into COS-1 cells. Full-length ACSVL1 and ACSVL3 were also each cloned into pcDNA3, and these constructs, as well as the empty vector, were each stably expressed in COS-1 cells for comparison.

**Figure 3.**
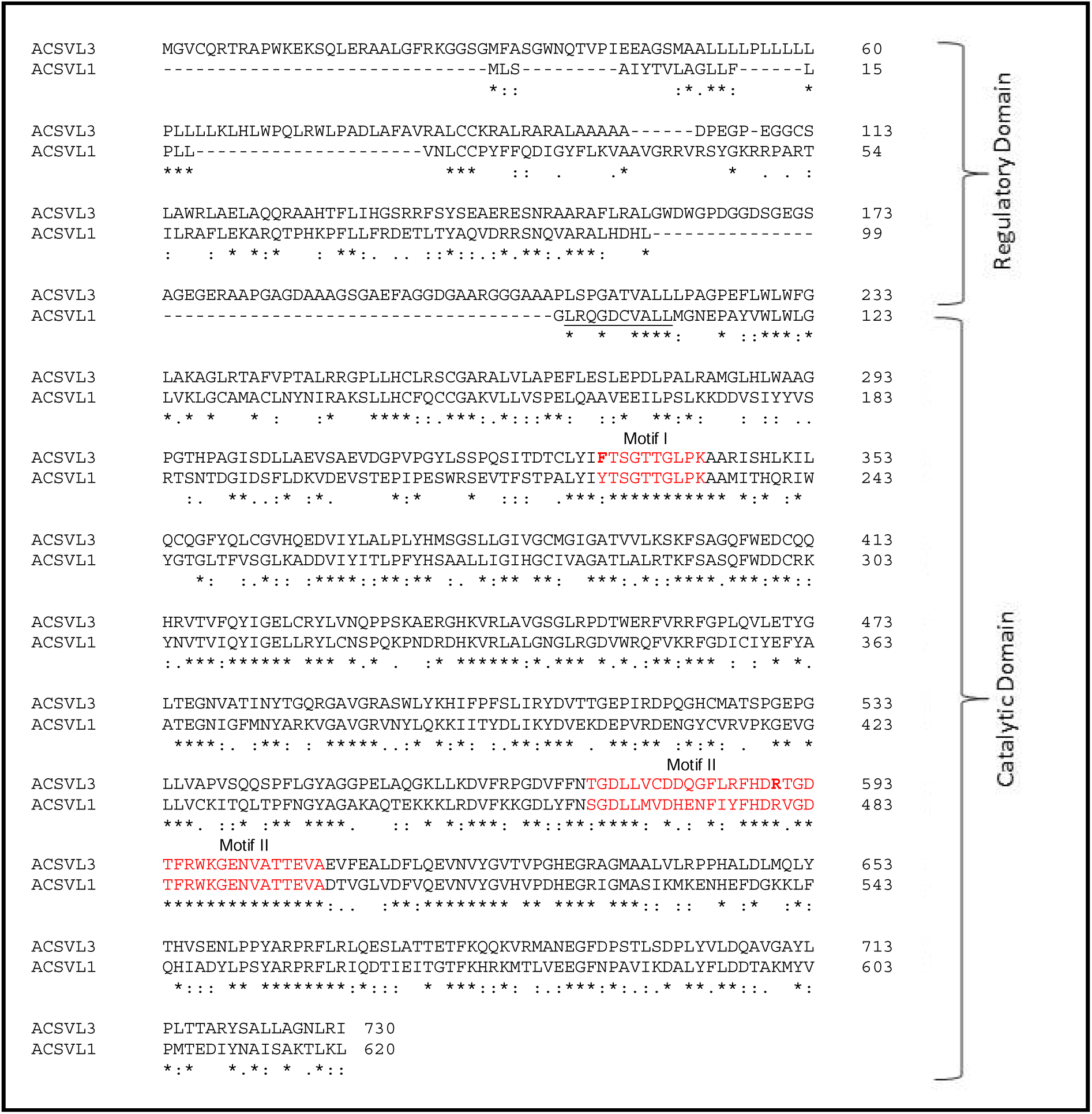
Sequence alignment of ACSVL3 and ACSVL1. The sequences of the homologous enzymes ACSVL3 (top row) and ACSVL1 (bottom row) were aligned using ClustalW at EMBL-EBI [Larkin et al., 2007; Goujon et al., 2010]. Regulatory and catalytic domains are denoted. The regulatory domain of ACSVL3 is 110 bp longer than the regulatory domain of ACSVL3. The start of the region of consistent homology, which is the proposed boundary between regulatory and catalytic domains, is underlined. The first leucine (L) in the underlined sequence marks the point of transition between the ACSVL1 N-terminus and the ACSVL3 C-terminus in the ACSVL1/3 chimera. The two regions in red typeface are conserved domains necessary for catalysis. The first region (Motif I) is an AMP biding domain, and the second (Motif II) is the proposed fatty acid binding “signature motif”. The phenylalanine residue at the start of Motif I and the arginine residue at the center of Motif II (bold red type) were changed to alanine residues using site-directed mutagenesis to create ACSVL1/3Mut1 and ACSVL1/3Mut2, respectively

**Figure 4.**
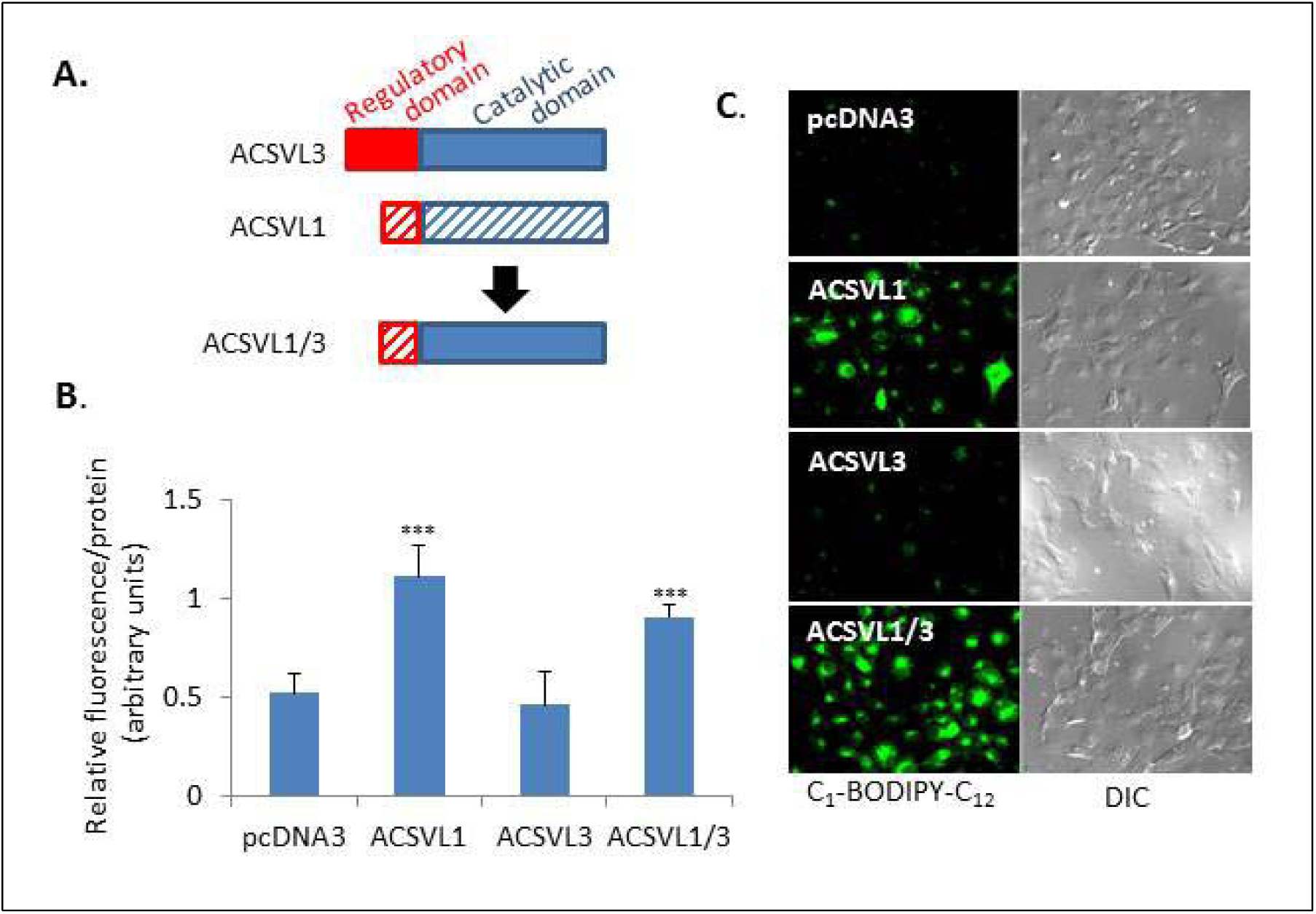
Characterization of the ACSVL1/3 hybrid enzyme. A. The domain swapped hybrid is illustrated. COS-1 cells were transfected with the empty vector (pcDNA3) or a plasmid containing ACSVL1, ACSVL3 or the hybrid ACSVL1/3. These cells were evaluated for their ability to uptake C_1_-BODIPY-C_12_ by quantification of fluorescence in a plate reader (B) or immunofluorescence (C). For quantification of fluorescence, triplicates of COS-1 cells expressing each construct were evaluated. ACSVL1- and ACSVl1/3-expressing cells both showed increased uptake of C_1_-BODIPY-C_12_ compared to vector only cells (p < 0.001), demonstrating that the domain swap conferred the ability to uptake C_1_-BODIPY-C_12_ to the catalytic domain of ACSVL3.

Fluorescence microscopy and a quantitative fluorescence-based plate-reader assay were used to compare the uptake of C_1_-BODIPY-C_12_ by COS-1 cells expressing ACSVL1, ACSVL3, the hybrid ACSVL1/3, and the empty vector pcDNA3. By microscopy, cells expressing ACSVL3 and pcDNA3 do not take up significant C_1_-BODIPY-C_12_ while ACSVL1 and ACSVL1/3 both take up this fluorescent FA analog at similar levels (Figure 4C). By the plate-reader assay, cells expressing ACSVL3 and pcDNA3 again show similar, lower levels of C_1_-BODIPY-C_12_ uptake, while cells expressing ACSVL1 and ACSVL1/3 exhibit similar, higher levels of uptake (Figure 4B).

### Mutations in the catalytic domain of ACVL1/3 hybrids are deficient in C_1_-BODIPY-C_12_ uptake

In the vectorial acylation model of cellular FA uptake (see Discussion), transport of the FA by an ACS is coupled to its enzyme activity [Black and DiRusso, 2003]. All 26 human and rodent ACS enzymes contain two highly conserved regions (Motif I and Motif II, shown in red typeface in Figure 4) that contain catalytically essential amino acids. To assess whether the ACS activity of ACSVL1/3 was essential for C_1_-BODIPY-C_12_ uptake, site-directed mutagenesis was used to change one catalytically critical residue in each of the two conserved regions of the chimeric protein. In the first mutant protein, ACSVL1/3mut1, the initial phenylalanine residue of the 10 amino acid Motif I was mutated to alanine; in the second, ACSVL1/3mut2, the centrally located arginine of Motif II was mutated to alanine. Mutation of either of these residues was previously shown to abolish ACS enzyme activity [Weimar JBC02; Black JBC97]. COS-1 cells transiently expressing ACSVL1/3, ACSVL1/3mut1, or ACSVL1/3mut2 were assessed for their ability to take up C_1_-BODIPY-C_12_. As shown in Fig. 5, ACSVL1/3 expressing cells readily took up the fluorescent fatty acid, while neither of the mutants promoted C_1_-BODIPY-C_12_ uptake. These data suggest that functional catalytic activity is required along with N-terminal domains for ACSVL1/3-mediated FA uptake.

**Figure 5.**
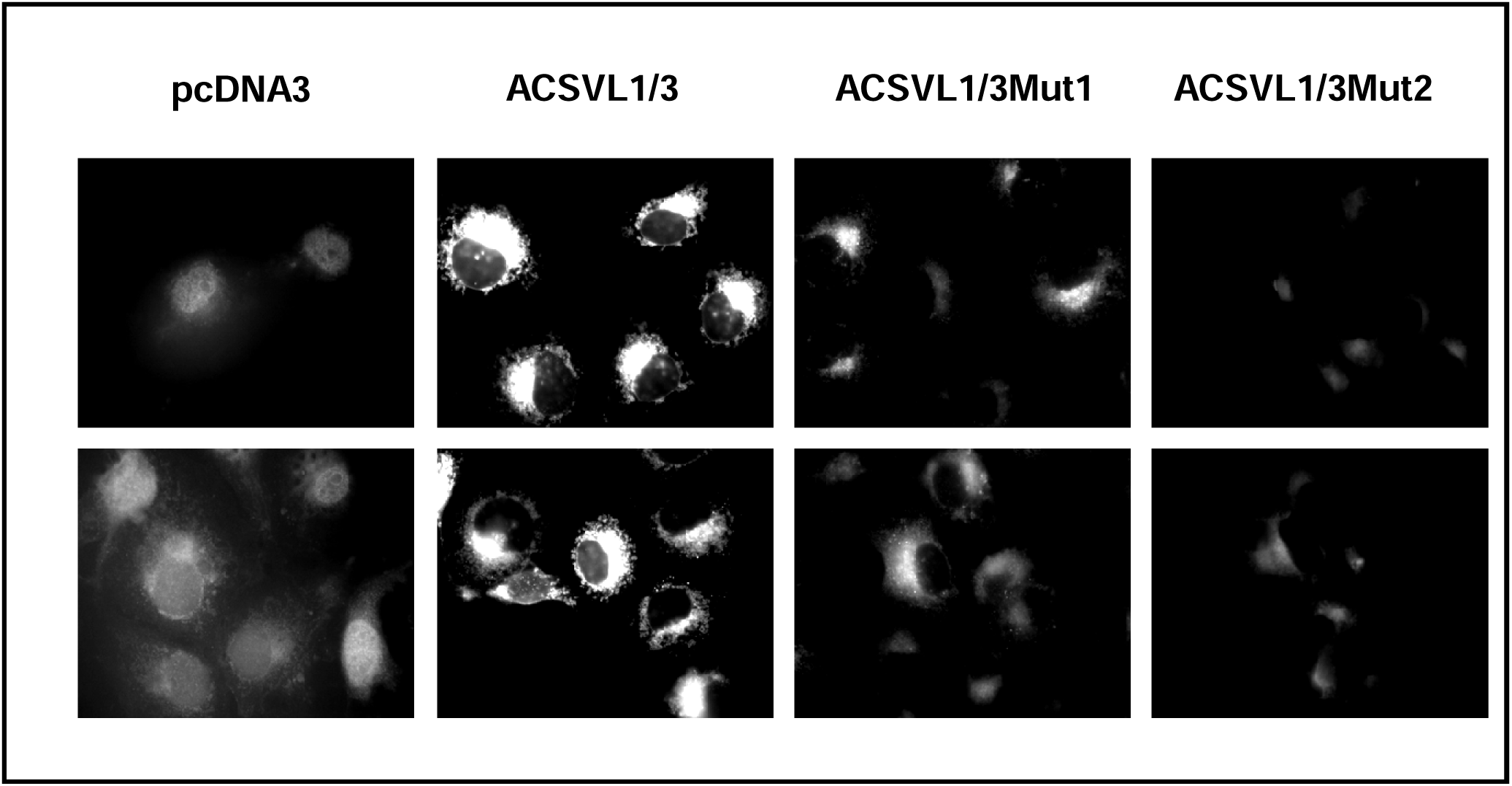
Uptake of C_1_-BODIPY-C_12_ by COS-1 cells expressing ACSVL1/3 catalytically deficient mutants. Site-directed mutagenesis was used to mutate catalytically critical amino acid residues of Motif I and Motif II in the ACSVL1/3 hybrid enzyme (see Fig. 2). The first amino acid of Motif I (phenylalanine) was changed to alanine to produce ACSVL1/3Mut1, and the central amino acid of Motif II (arginine) was changed to alanine to produce ACSVL1/3Mut2. COS-1 cells stably expressing ACSVL1/3, ACSVL1/3Mut1, ACSVL1/3Mut2 or harboring the empty pcDNA3 vector were assessed for their ability to take up the fluorescent FA analog C_1_-BODIPY-C_12_. Both mutants showed a severe deficiency in their ability to take up C_1_-BODIPY-C_12_.

### Most ACSVL1 inhibitors also inhibit the acyl-CoA synthetase activity of ACSVL1/3 and its ability to promote C_1_-BODIPY-C_12_ uptake

The same set of ACSVL1 inhibitors that were screened for their ability to decrease stearoyl-CoA synthetase activity assay (Figure 2) were tested to see if they would inhibit uptake of C_1_-BODIPY-C_12_ in COS-1 cells expressing ACSVL1/3. Each was added in multiple concentrations from 0.1 to 5 μM in duplicate wells of a 96-well plate. Many of the inhibitors showed a dose-dependent inhibition of C_1_-BODIPY-C_12_ uptake (Figure 6). Again, the compounds CB5 and CB16.2 were found to be the most potent of these candidate ACSVL3 inhibitors.

**Figure 6.**
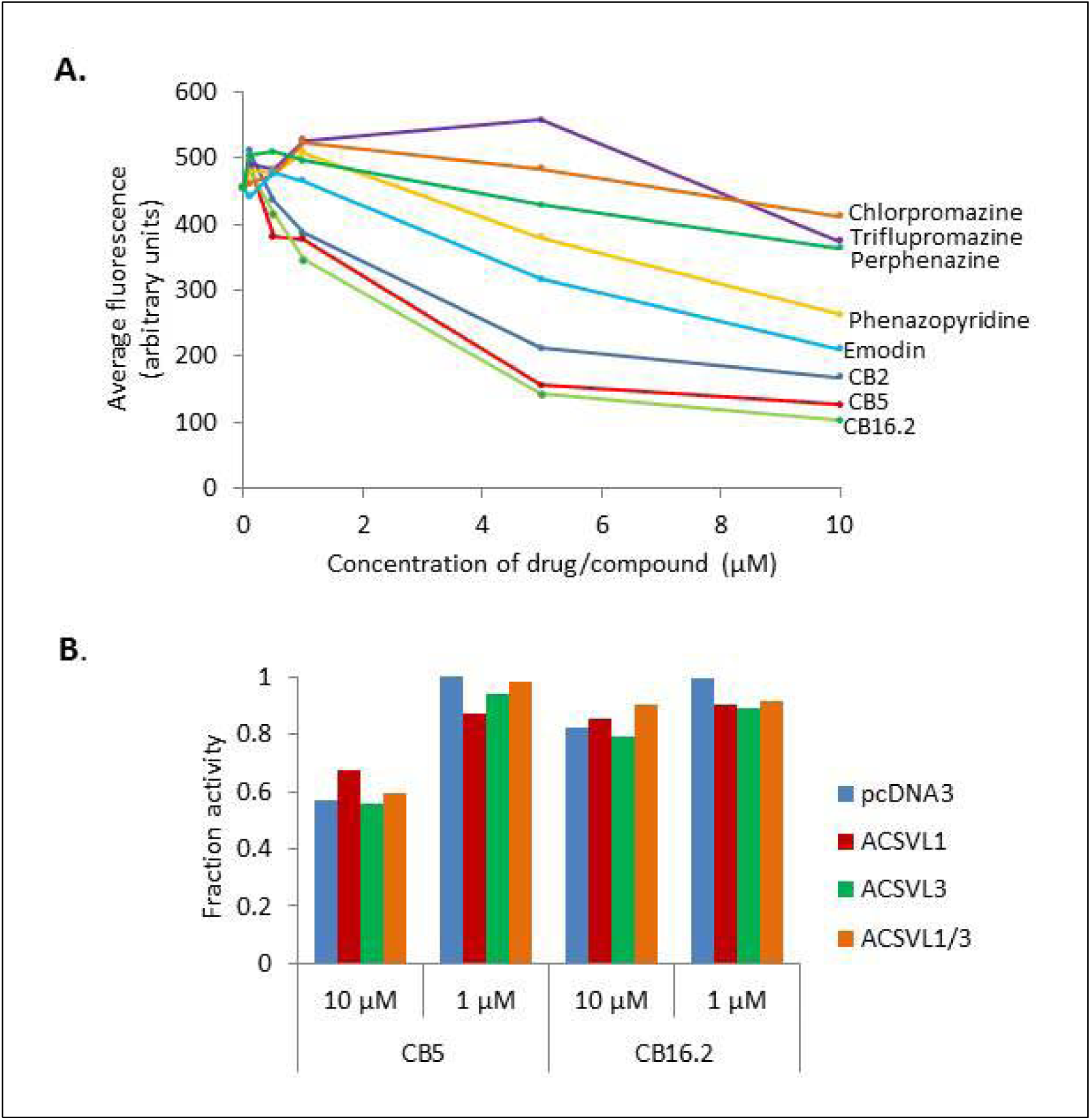
Effect of inhibitors on C_1_-BODIPY-C_12_ uptake and ACS activity by COS-1 cells expressing ACSVL1/3. A. Fluorescent FA uptake. COS-1 cells expressing ACSVL1/3 were treated with various concentrations of ACSVL1 inhibitors as described in Methods. Uptake of fluorescent C_1_-BODIPY-C_12_ was quantified using a SpectraMax M5 microplate reader and average fluorescence plotted. Most drugs and compounds inhibited C_1_-BODIPY-C_12_ uptake in a dose-dependent manner. B. ACS activity. COS-1 cells expressing ACSVL1, ACSVL3, ACSVL1/3, or the empty vector pcDNA3 were assayed for their ability to activate [1-^14^C]stearic acid (C18:0) to its CoA derivative in the absence or presence of CB5 and CB16.2. The fraction of activity in drug-treated vs. untreated (DMSO only) cells is shown. 10 µM CB5 inhibited Cos-1 cells containing each vector, including empty vector pcDNA3, most likely due to inhibition of other ACS enzymes endogenousl expressed by COS-1 cells.

CB5 and CB16.2 were added to acyl-CoA synthetase assays measuring the conversion of radiolabeled stearic acid to stearyl-CoA to assess if they would have a similar inhibitory effect on ACSVL1/3 as they did on ACSVL3. Both drugs were tested at concentrations of 10 μM and 1 μM. Pellets from COS-1 cells overexpressing each construct (ACSVL1, ACSVL3, ACSVL1/3, and pcDNA3) were assayed. While 10 μM and 1 μM CB16.2 as well as 1 μM CB5 had minor inhibitory effects, 10 μM CB5 inhibited stearoyl-CoA synthetase activity of all constructs (Figure 6B and 6C). The inhibition seen in cells expressing pcDNA3 can be attributed to inhibition of other long- and very long-chain ACSs endogenously expressed in COS-1 cells.

### Pilot screening study of >2150 library compounds identified additional candidate ACSVL3 inhibitors

Using the approach outlined in Figure 6, we conducted a pilot study to determine whether we could identify additional compounds that might be candidate ACSVL3 inhibitors. Libraries screened include subsets of Johns Hopkins Clinical Compound Library 20 and the ChemBridge Corporation CNS-set. Of the ∼1280 compounds from the Johns Hopkins library that we tested, 32 compounds yielded a decrease in fluorescence that was more than 3 standard deviations lower than no drug. Similarly, of ∼880 compounds from the ChemBridge library tested, 27 lowered C_1_-BODIPY-C_12_ uptake by the same criteria. There are several validation studies that must be carried out before considering any of these “hits” as ACSVL3 inhibitors, including a repeat screen. Compounds that reproducibly lower C_1_-BODIPY-C_12_ uptake must also be tested for fluorescence quenching, which would artificially decrease fluorescence, and to ensure that they do not cause cell permeabilzation, which would dilute any C_1_-BODIPY-C_12_ taken up by the chimeric cells. Nonetheless, these studies suggest that COS-1 cells expressing the hybrid ACSVL1/3 construct could form the basis of a high-throughput screen for ACSVL3 inhibitors – compounds that ultimately may prove clinically effective in treating glioma and other human malignancies.

## DISCUSSION

In this study, we report results of our efforts to identify inhibitors of ACSVL3, one of six members of the very long-chain acyl-CoA synthetase family [Watkins 2008; Watkins et al., 2007]. Hypothesizing that inhibitors of a structurally and functionally related enzyme such as ACSVL1 might also inhibit ACSVL3, we tested several compounds and identified several that inhibit the activity of both enzymes, but to different degrees. We also developed a 96 well-based assay that could form the basis for conducting an enzyme-specific high-throughput screen of chemical libraries for ACSVL3 inhibitors.

There were two main reasons for searching for an ACSVL3 inhibitor. First, a specific inhibitor would be a valuable tool to help understand the biology of ACSVL3 and tumorigenesis, and second, an inhibitor could have therapeutic potential in patients suffering from glioma. These experiments have identified inhibitors of ACSVL3 activity, and the most robust one, CB5, shows potential for both purposes.

One difficulty in identifying specific inhibitors of acyl-CoA synthetases is that many different ACSs with overlapping function and substrate specificity can be normally present in the same cell type (Table 2). In vivo, this can be explained by the observation that different enzymes with similar activities reside in distinct subcellular compartments. U87MG cells contain other long-chain and very-long chain ACS family members that are predicted to activate stearic acid to stearyl-CoA, making it difficult to create a highly specific screening assay for inhibition of ACSVL3 enzyme activity alone. Some knowledge of specificity was obtained by comparing ACS activity in U87MG cell suspensions to that in U87-KO cell suspensions. If the inhibitors screened are somewhat specific for ACSVL3, they are predicted to inhibit stearic acid activation more robustly in U87MG than in U87-KO cellular protein. An ideal inhibitor would be one that displayed no significant inhibition of other ACS family enzymes, and would inhibit stearic acid activation in U87MG cells but not in U87-KO cells. However, due to the high degree of homology in the catalytically critical regions of ACS enzymes, it is unlikely that a completely specific inhibitor will easily be found.

The U87-KO line has been very useful in characterizing the role that ACSVL3 plays in lipid metabolism in glioma, and halting ACSVL3 activity with an inhibitor would in theory produce the same phenotypes exhibited in the U87-KO line. For therapeutic uses of identified inhibitors, complete specificity may not be crucial, as chemotherapeutic agents are used more as acute treatments than as chronic treatments. The primary consideration is that the inhibitor abrogates cancerous phenotypes without significant side-effects.

Based on their studies of bacterial FA transport, Black and DiRusso proposed the concept of vectorial acylation in which transport of a FA across the cell membrane was “pulled” by activation of the transported FA to its CoA derivative [DiRusso and Black, 2004]. In some cases, separate transport proteins and the ACSs form a complex to facilitate FA uptake, while in others, the two processes are carried out by a single integral membrane protein [reviewed in Arias-Barrau, DiRusso and Black, 2009]. Three members of the FATP/ACSVL family (FATP1/ACSVL4, FATP2/ACSVL1 and FATP4/ACSVL5) are capable of carrying out both processes, and Black and DiRusso exploited the fatty acid transport property of ACSVL1 to develop a high-throughput screen for inhibitors of this enzyme [Black and DiRusso, U.S. Patent No. 7,070,944, 2006]. They produced a strain of yeast deficient in both fatty acid transport and acyl-CoA synthetase activity, and exogenously expressed human ACSVL1. ACSVL1-overexpressing cells took up the fluorescent long chain fatty acid analog C_1_-BODIPY-C_12_ while those lacking ACSVL1 did not. Using fatty acid uptake as a surrogate for acyl-CoA synthetase activity, over 100,000 drugs and compounds were screened [Li et al., 2007; Sandoval et al., 2010]. Yeast cells were treated with compounds from three libraries: the SpectrumPlus library from MicroSource Discovery Systems Inc., the NCI Diversity Set Compound Library and the ChemBridge Corporation compound library. The screen identified several compounds that inhibited the uptake of C_1_-BODIPY-C_12_. The SpectrumPlus screen revealed 28 ACSVL1 potential inhibitors, many of which were tricyclic, phenothiazine-derived antipsychotic drugs. The Diversity Set/ChemBridge screen revealed 216 potential ACSVL1 inhibitors, which clustered into 5 structural classes. The C-terminal catalytic domains of ACSVL1 and ACSVL3 are highly homologous (Figure 2). These domains bind both fatty acids and ATP and therefore are responsible for the conversion of long- and very-long chain fatty acids to their acyl-CoA derivatives. However, the smaller, N-terminal domains of ACSVL1 and ACSVL3 are significantly different. While the exact purpose of the N-terminal domains is not completely understood, it is assumed to be a regulatory domain, possibly involved in subcellular location determination, membrane association or fatty acid transport. Since ACSVL1 is homologous to ACSVL3 throughout their catalytic domains, we expected that ACSVL1 inhibitors might also inhibit the ACS enzyme activity of ACSVL3.

In this study, we chose representative ACSVL1 inhibitors to test for inhibition of ACSVL3. We used two metrics to characterize potential inhibitors. First was the inhibition of the thioesterification of [^14^C]stearic acid to [^14^C]stearoyl-CoA. Stearic acid was chosen above other long- and very long-chain fatty acids because it is activated preferentially by ACSVL3 at a relatively high rate. The second metric was the inhibition of uptake of C_1_-BODIPY-C_12_ in COS-1 cells expressing the chimeric protein ACSVL1/3. By both metrics, CB5 was the most robust inhibitor of ACSVL3 activity of the compounds tested. Studies to assess how CB5 affects the malignant behavior of U87MG cells both in vitro and in mouse xenografts are currently underway.

As a potent inhibitor of ACSVL1 [Sandoval et al., 2010], we expected that CB16.2 might also be a good inhibitor of ACSVL3. However, results were mixed. CB16.2 inhibited uptake of C_1_-BODIPY-C_12_ in COS-1 cells expressing ACSVL1/3 nearly as well as CB5. At a concentration of 80 μM, CB16.2 inhibited conversion of stearate to stearyl-CoA less robustly than CB5 but better than any other inhibitor, but at 10 μM, CB5 significantly outperformed CB16.2. These inconsistencies may be due to the fact that CB16.2 appeared to be less soluble than the other inhibitors, including CB5, in the solvent used. It seemed more likely to precipitate out of DMSO, and when the stock DMSO solution was added to the aqueous assays, it appeared to form a suspension, which may or may not have been stable. CB16.2 may have been a better inhibitor by our metrics if it was delivered in a different solvent.

While all compounds listed in Table 1 showed some degree of ACSVL3 inhibition, not all ACSVL1 inhibitors also inhibit ACSVL3. We previously conducted preliminary studies in which compounds were tested for potential inhibition of ACSVL3 activity using a suboptimal ACS substrate, [^14^C]palmitic acid. In both the preliminary study and the present report, ACSVL1 inhibitors triflupromazine, phenazopyridine, chlorpromazine, emodin and perphenazine were inhibitory to ACSVL3. In contrast, several other compounds reported to inhibit ACSVL1 such as clomipramine, clozapine, promazine and cetrimonium bromide [Li et al., 2007] did not appear to be inhibitory toward ACSVL3 (data not shown). Thus, not all ACSVL1 inhibitors would be expected to inhibit ACSVL3.

Unlike ACSVL1, ACSVL3 does not promote FA uptake, preventing us from developing a screening tool for ACSVL1 inhibitors using the yeast-based platform developed by Black, DiRusso and colleagues described above. Hypothesizing that the N-terminus of ACSVL1 might be necessary for the FA transport function of this enzyme, chimeric ACSVL1/3 was created to couple the putative transport capability of the ACSVL1 N-terminus to the enzymatic activity of ACSVL3 (Figures 4 and 5). Our hypothesis proved to be correct and suggests that both the N-terminal transport function and the C-terminal catalytic domains are necessary for FA uptake.

Using ACSVL1/3-expressing COS-1 cells, we developed a 96-well plate fluorescence-based assay to compare several potential ACSVL3 inhibitors at multiple concentrations (Figure 6). This assay was also used to establish proof-of-concept that it could be used to screen chemical libraries to identify additional inhibitors of ACSVL3. Subsets of a library of approved drugs and natural products, and of the ChemBridge CNS-Set library, were screened for reduction of C_1_-BODIPY-C_12_ uptake by measuring intracellular fluorescence using a plate reader. Out of more than 2000 compounds tested, a first-pass screen detected more than 50 compounds that lowered intracellular fluorescence by at least 3 standard deviations from the mean of the control (no drug added) signal. We expect that most of these will not withstand further scrutiny, primarily due to C_1_-BODIPY-C_12_ signal quenching, toxicity, or cellular membrane disruption. Drugs that remain as candidate inhibitors will then be tested for inhibition of ACSVL3 synthetase activity in stably transfected COS-1 cells and U87 cells, with the ultimate goal of finding one or more highly specific ACSVL3 inhibitors.

In conclusion, this report suggests that inhibitors of ACSVL3 exist. If one or more specific ACSVL3 inhibitors can be identified, they may have therapeutic potential in treating glioma. Further studies testing the potential of the compound CB5 in abrogating the malignant properties of U87MG cells both in vitro and in mice are in progress.

## ACKNOWLEDGEMENTS

The authors thank Drs. Paul Black and Concetta DiRusso (Dept. of Biochemistry, Univ. of Nebraska, Lincoln NE) for providing several ACSVL1 inhibitors used in this study. The authors thank Dr. Jun O. Liu (Department of Pharmacology and Experimental Therapeutics, Johns Hopkins University School of Medicine, Baltimore MD) for making the Johns Hopkins Clinical Compound Library available for this study. We also thank Drs. Akhilesh Pandey and Raja Sekhar Nirujogi (McKusick-Nathans Institute of Genetic Medicine, Johns Hopkins University School of Medicine, Baltimore MD) for proteomic analysis of human astrocytes and U87MG cells. This work was supported by NIH grant NS062043.

